# The role of islet lipid composition remodeling in regulation of beta-cell death via ADP-ribosyl-acceptor glycohydrolase ARH3 signaling in insulitis

**DOI:** 10.1101/2020.03.23.004481

**Authors:** Ernesto S. Nakayasu, Cailin Deiter, Jennifer E. Kyle, Michelle A. Guney, Dylan Sarbaugh, Ruichuan Yin, Yi Cui, Carrie D. Nicora, Farooq Syed, Jonas Juan-Mateu, Raghavendra G. Mirmira, Carmella Evans-Molina, Decio L. Eizirik, Bobbie-Jo M. Webb-Robertson, Kristin Burnum-Johnson, Galya Orr, Julia Laskin, Thomas O. Metz, Lori Sussel, Charles Ansong

**Affiliations:** Biological Sciences Division, Pacific Northwest National Laboratory, Richland, WA, USA; Barbara Davis Center for Diabetes, University of Colorado Anschutz Medical Center, Aurora, CO, USA; Department of Chemistry, Purdue University, West Lafayette, IN, USA; Environmental and Molecular Sciences Laboratory, Pacific Northwest National Laboratory, Richland, WA, USA; Media Lab, Massachusetts Institute of Technology, Cambridge, MA, USA; Center for Diabetes and Metabolic Diseases and the Herman B Wells Center for Pediatric Research, Indiana University School of Medicine, Indianapolis, IN, USA; ULB Center for Diabetes Research, Université Libre de Bruxelles (ULB), Brussels, Belgium; Centre for Genomic Regulation (CRG), The Barcelona Institute of Science and Technology, Barcelona, Spain; Kovler Diabetes Center and Department of Medicine, The University of Chicago, Chicago, IL, USA; Indiana Biosciences Research Institute Diabetes Center, Indianapolis, IN, USA; Department of Biostatistics and Informatics, University of Colorado Anschutz Medical Center, Aurora, CO, USA

**Keywords:** Type 1 diabetes, insulitis, lipidomics, phospholipase, poly(ADP)ribosylation, β-cell death

## Abstract

Lipids have been implicated as mediators of insulitis and β-cell death in type 1 diabetes development, but the mechanisms underlying this association are poorly understood. Here, we investigated the changes in islet/β-cell lipid composition using three models of insulitis: human islets and EndoC-βH1 β-cells treated with the cytokines IL-1β and IFN-γ, and islets from non-obese diabetic mice. Across all three models, lipidomic analyses showed a consistent change in abundance of the lysophosphatidylcholine, phosphatidylcholine and triacylglycerol species. We also showed that lysophosphatidylcholine and its biosynthetic enzyme PLA2G6 are enriched in murine islets. We determined that the ADP-ribosyl-acceptor glycohydrolase ARH3 is regulated by cytokines downstream of PLA2G6, which in turn regulates proteins involved in apoptosis, lipid metabolism, antigen processing and presentation and chemokines. ARH3 reduced cytokine-induced apoptosis, which may represent a negative feedback mechanism. Overall, these data show the importance of lipid metabolism in regulating β-cell death in type 1 diabetes.

**Highlights:**

- Lipidomics of 3 insulitis models revealed commonly regulated lipid classes.
- Identification of 35 proteins regulated by cytokines via PLA2G6 signaling.
- ARH3 reduces cytokine-induced apoptosis via PLA2G6 regulation.
- ARH3 regulates the levels of proteins related to insulitis and type 1 diabetes.

## INTRODUCTION

Type 1 diabetes (T1D) affects over 1.25 million people in the U.S. and is characterized by the autoimmune destruction of the insulin-producing β cells of the pancreas (DiMeglio et al., 2018). This results in the disruption of blood glucose regulation, the development of microvascular and macrovascular complications, and ultimately reduces life expectancy of individuals with T1D by approximately 12 years (Huo et al., 2016). Currently, the treatment of T1D relies on exogenous insulin administration and there is no cure or permanent remission from the disease. Therefore, there is an urgent need to better understand the mechanisms that contribute to β-cell loss in order to inform the development of novel treatment therapies.

Lipids play major roles in all tissues of the body, being the major structural component of membranes, energy storage molecules and cell-signaling mediators. T1D is associated with changes in serum lipid profiles after disease onset, including increases in blood triacylglycerol and cholesterol levels, which are often associated with poor glycemic control (Verges, 2009). However, limited information is available regarding the roles of lipids in contributing to disease pathogenesis. For instance, the Oresic group found that circulating lipids function as biomarkers for autoimmunity and T1D progression (Lamichhane et al., 2018; Oresic et al., 2013). In terms of cell signaling, lipids have been shown to be important mediators of the islet inflammatory process (insulitis) (Bone et al., 2015; Dobrian et al., 2019; Syed et al., 2019). During insulitis, pro-inflammatory cytokines activate phospholipases, such as inducible phospholipase A2 (PLA2), leading to the cleavage of phosphatidylcholine (PC) into lysophosphatidylcholine (LPC) (Barbour et al., 2015; Bone et al., 2015; Lei et al., 2014). This process also leads to the release of fatty acids, such as arachidonic acid, which can be converted into prostaglandins, leukotrienes and thromboxanes that activate pro-apoptotic signaling and contribute to β-cell death (Chambers et al., 2007; Ma et al., 1996).

Currently, the role of the islet lipidome during the progression of T1D is poorly understood. Mass spectrometry analysis of islets has identified LPC and ceramides as important molecules in β-cell death (Lei et al., 2014). However, the lipidomics coverage in that study was limited and recent advances in technology can provide a more comprehensive coverage of the islet lipidome. Here, we used 3 models of insulitis and type 1 diabetes progression to study the changes in islet/β cell lipid composition during this process. Unbiased lipidomics analyses were performed in 1) human islets and 2) a human β-cell line exposed to the proinflammatory cytokines IL-1β and IFN-γ, that induce a molecular signature similar to the one observed in β cells from patients affected by type 1 diabetes (Eizirik et al., 2020), and 3) in islets from non-obese diabetic (NOD) mice. The analysis showed a consistent regulation of LPC, PC with long unsaturated fatty acid chains and triacylglycerol species across the different models. To further investigate LPC regulation, we performed mass spectrometry imaging analysis and microscopy of its biosynthetic enzyme PLA2G6 in mouse islets. The results showed that both LPC and PLA2G6 are enriched in islets. We also investigated the signaling mediated by PLA2G6 in mouse Min6 β cells using RNAi followed by proteomics analysis. This analysis determined that the ADP-ribosyl-acceptor glycohydrolase ARH3 is regulated by cytokines and, based on RNAi experiments targeting ARH3, we observed that this protein provides a complex negative feedback mechanism that reduces cytokine-induced apoptosis.

## MATERIAL AND METHODS

### Mice

Female 6-week-old non-obese diabetic NOD/ShiLtJ (#001976) and non-obese diabetes-resistant NOR/LtJ (#002050) mice were purchased from Jackson Labs. Husbandry of mice and experimental procedures were performed according an approved IACUC protocol at the University of Colorado (#00045). Mice were housed for < 1 week prior to islet isolation. Approximately 320-520 islets (n=3) were isolated using Histopaque centrifugation gradient and further handpicking of islets as described previously (Beilke et al., 2005).

### Human pancreatic islet and EndoC-βH1 cells

The lipidomics data of the human islets (n=10) (**Supplementary Table 1**) and EndoC-βH1 cells (n=3) treated with 50 U/mL IL-1β + 1000 U/mL IFN-γ were collected from the same samples of proteomic analyses carried out by us in (Nakayasu et al., 2020a) and (Ramos-Rodriguez et al., 2019), respectively. Culture maintenance, treatment and harvesting information are described in detail elsewhere (Nakayasu et al., 2020a; Ramos-Rodriguez et al., 2019).

### MIN6 β cell line culture and treatment

MIN6 cells were cultured in DMEM containing 10% FBS and 1% penicillin-streptomycin and maintained at 37ºC in 5% CO_2_ atmosphere. For knockdown experiments, cells were transfected using Lipofectamine RNAiMAX (Invitrogen) with SMARTpool ON-TARGETplus non-targeting scramble siRNA (control) or siRNA targeting either *PLA2g6* or *ARH3* (Dharmacon). In order to achieve robust knockdown of ARH3, cells were transfected a second time with scramble or ARH3 siRNA 24 h after the first transfection. In experiments utilizing cytokines, cells were treated for 24 h with 100 ng/mL IFN-γ, 10 ng/mL TNF-α, and 5 ng/mL IL-1β.

### Lipidomic analysis

Samples were subjected to metabolite, protein and lipid extraction (MPLEx) (Nakayasu et al., 2016) with all the procedure done on ice or 4 ºC to reduce sample degradation. Cells and islets were suspended in water and extracted with 5 volumes of cold (−20 °C) chloroform/methanol solution (2/1, v/v) by incubating 5 min on ice and vigorously vortexing. Samples were centrifuged for 10 min at 16,000xg at 4°C to separate different liquid phases. Solvent layers containing metabolites and lipids were collected into glass autosampler vials and dried in a vacuum centrifuge. Protein pellets were washed by adding 1 mL of cold (−20°C) methanol and centrifuging 10 min at 16,000xg at 4°C. The supernatant was discarded, and pellets were dried in a vacuum centrifuge before digesting with trypsin.

Extracted lipids were resuspended in methanol and loaded on a reverse phase column connected to a NanoAcquity UPLC system (Waters) and interfaced with a Velos Orbitrap mass spectrometer (Thermo Fisher) (Dautel et al., 2017) (see **Supplementary Table 2** for parameters). Lipid species were identified using LIQUID (Dautel et al., 2017) and manually validated based on the retention time, precursor isotopic profile, diagnostic fragments from head groups and fatty acyl chains. Isomers were named in alphabetical order based on their elution times. The features of the identified lipid species were extracted and aligned using targeted peak detection in MZmine (Pluskal et al., 2010). Peak intensity tolerance was set at 20% with a noise level of 5e3, while the mass and retention time tolerances were 0.008 m/z and 0.3 min, respectively. Chromatograms were built with a time span of 0.1 min, minimum height of 5e3 and mass tolerance of 0.008 m/z. Peaks were then aligned with joint aligner with mass tolerance of 0.01 m/z, retention time tolerance of 0.2 min, and weights of both retention time and m/z of 1. Missing values were filled with gap filling based on the same retention time and mass tolerance of 0.008 m/z. The lipidomics data were log2 transformed and subjected to quality control analysis to identify any sample issues via Principal Component Analysis (PCA), robust to missing data for proteomics (3), as well as a robust PCA approach based on the distributional properties of the measured biomolecules for each sample (4). No sample-level issues were detected and subsequently the lipidomics data were evaluated for total abundance bias and, with none found, were global median centered (5). Given the experimental design, statistics were then performed via a standard paired t-test followed by a multiple test Bonferroni correction (6).

### Proteomic analysis

Proteins were dissolved in 50 mM NH_4_HCO_3_ containing 8 M urea and 10 mM dithiothreitol. After incubating for 1 h at 37 ºC with shaking at 800 rpm, 400 mM iodoacetamide was added to a final concentration of 40 mM, and the mixture incubated for another hour in the dark at room temperature. The reaction mixture was 8-folds diluted with 50 mM NH_4_HCO_3_, and 1 M CaCl_2_ was added to a final concentration of 1 mM. Proteins were digested for 3 h at 37 ºC using trypsin at 1:50 enzyme:protein ratio. Digested peptides were desalted by solid-phase extraction using C18 cartridges (Discovery, 50 mg, Sulpelco) and dried in a vacuum centrifuge.

Peptides were analyzed on a Waters NanoAquity UPLC system coupled with a Q-Exactive mass spectrometer (see **Supplementary Table 2** for parameters). Data were processed with MaxQuant software (v.1.5.5.1 and v1.6.14.0) (Cox and Mann, 2008) by matching against the mouse reference proteome database from Uniprot Knowledge Base (downloaded on August 14, 2018 and on August 31, 2020). Searching parameters included protein N-terminal acetylation and oxidation of methionine as variable modifications, and carbamidomethylation of cysteine residues as fixed modification. Mass shift tolerance was used as the default setting of the software. Only fully tryptic digested peptides were considered, allowing up to two missed cleaved sites per peptide. Quantification of proteins was done using the intensity-based absolute quantification (iBAQ) method (Schwanhausser et al., 2011). Data was log2 transformed, and normalized by linear regression and central tendency using InfernoRDN (former Dante) (Polpitiya et al., 2008). Statistically significant proteins were determined by ANOVA or by *t*-test.

### Bioinformatics analysis

The significantly different lipids and proteins were submitted to ontology/function-enrichment analysis using Lipid MiniOn (Clair et al., 2019) and DAVID (Huang da et al., 2009) tools, respectively. For the Lipid MiniOn analysis, the full list of identified lipids was set as the background and the significantly different species as the query. Ontologies were considered enriched with a p ≤ 0.05 using the Fisher’s exact test. For the DAVID analysis, the differentially abundant proteins were set as the query and the entire genome was set as the background. Only enriched pathways (p ≤ 0.05) of the KEGG database were used and they were grouped based on shared proteins using Enrichment Map (Merico et al., 2010).

### Tissue processing and immunohistochemistry

Pancreata were dissected, washed in ice cold PBS, and fixed for 4 hours in 4% PFA at 4 °C. The tissue was then incubated in 30% sucrose overnight and frozen in OCT. 10 μm cryosections were taken and placed on Superfrost Plus slides (Fisher Scientific). Immunohistochemistry for PLA2G6 was performed using the Mouse on Mouse kit (Vector Labs) and the Vectastain ABC HRP kit (Vector Labs).

### Mass spectrometry imaging analysis

Nanospray desorption electrospray ionization (nano-DESI) mass spectrometry imaging was performed on a Q Exactive HF-X mass spectrometer (Thermo Fisher Scientific) (Yin et al., 2019). High-resolution nano-DESI probes were assembled using two fused silica capillaries pulled to O.D. 15-25 μm. A shear force probe with a tip diameter of ~10 μm was integrated with the nano-DESI probe and was used to precisely control the distance between the probe and the sample. The position of the samples was controlled by a motorized XYZ stage. Samples were scanned at a rate of 10 μm/s under the nano-DESI probe in lines with a step of 20 μm between the lines. A 9/1 (v/v) methanol/water mixture containing 200 nM LPC 19:0 (internal standard) was propelled through the nano-DESI probe at 500 nL/min (see **Supplementary Table 2** for parameters). A custom-designed software, MSI QuickView, was used for data visualization and processing. Ion images were generated by normalizing the signal of the analyte to the signal of the internal standard (LPC 19:0). Lipids were identified by matching based on the high mass accuracy against the species characterized in the lipidomics analysis.

### Fluorescence in-situ hybridization (FISH)

FISH experiments were performed as previously described (Cui et al., 2018). Ten oligonucleotide probes containing targeting and 3’ readout overhang domains were designed against the *Pla2g6* transcript coding region (see **Supplementary Table 3** for sequences). Each targeting domain was 20-nt long and complementary to the *Pla2g6* mRNA, with CG content of 40-60%, no self-repeats and inner loop structures. A secondary probe labeled with two Alexa647 molecules was used to hybridize with the 3’ overhang domain. Samples were fixed with fresh 4% paraformaldehyde. After quenching the residual paraformaldehyde was with 0.1% sodium borohydride, samples were permeabilized with 0.2% Triton-X 100 and stored in 70% ethanol. The primary and secondary probes (50 nM final concentration) and anti-insulin antibody (**Supplementary Table 3**) (1000-fold dilution) were diluted in hybridization solution (10% dextran sulfate, 15% formamide, 1× SSC, 3.4 mg/mL tRNA, 0.2 mg/mL RNase-free BSA, 2 mM ribonucleoside vanadyl complex) and incubated with the samples 37°C overnight in a humid chamber. Samples were rinsed with 15% formamide in 1× SSC, followed by staining with Atto 488-conjugated secondary antibody (**Supplementary Table 3**) and DAPI. Images were collected on an Olympus IX71-based single-molecule microscope equipped with 405 nm, 488 nm, and 640 nm solid lasers. Images were captured using a 100× oil immersion objective lens (NA 1.4) and an EMCCD camera (Andor iXon Ultra 897). Fluctuation localization imaging-based FISH (fliFISH) was used to extract the true location of *Pla2g6* mRNA in tissue sections to account for the background noise (Cui et al., 2018). The location for each *Pla2g6* transcript was so calculated based on the center of mass. Photoswitching was activated by using GLOX-containing buffer: 50 mM Tris, 10 mM NaCl, 10% glucose, 560 μg/mL glucose oxidase, 34 μg/mL catalase and 1% β-mercaptoethanol.

### Quantitative real-time PCR analysis

Cells were harvested and total mRNA was extracted using the RNeasy Mini kit (Qiagen). cDNA synthesis was performed using the iScript synthesis kit (Biorad) and qRT-PCR assays were performed using SsoAdvanced Universal SYBR Green Supermix. Expression levels were normalized to the housekeeping gene GAPDH and quantified using the delta CT method (see **Supplementary Table 3** for oligonucleotide information).

### Western Blotting

Cells were harvested in RIPA buffer containing protease inhibitors and proteins were electrophoresed and transferred to PVDF membranes. After blocking in 5% milk in TBS containing 0.1% Tween 20, membranes were incubated in specific primary antibodies at 4°C overnight. Anti-mouse and anti-rabbit horseradish peroxidase-conjugated antibodies (**Supplementary Table 3**) were used for secondary antibodies and enhanced chemiluminescent substrate was used for signal detection.

## RESULTS

### Lipidome analysis of β-cell death/type 1 diabetes models

We studied three models of insulitis and T1D progression: (I) Human EndoC-βH1 cells exposed to the cytokines IL-1β + INF-γ for 48 h, (II) human islets exposed to the same cytokines for 24 h, and (III) islets from non-obese diabetic (NOD) mice at the pre-diabetic stage (6-week old) vs. age-matched NOR mice. To verify that under the present experimental conditions 6 weeks corresponds to the initial T1D developmental stages of NOD mice, we performed proteomics analysis of islets from both NOD and NOR mice (**Supplementary Table 4-5**) and compared the results against published proteomics data of EndoC-βH1 cells and human islets exposed IL-1β + INF-γ (Nakayasu et al., 2020a; Ramos-Rodriguez et al., 2019). We observed an upregulation of inflammatory markers, such as the antigen transport protein Tap1, the transcription factor Stat1 and the interferon-induced guanylate-binding protein GBP2 (**Supplementary Figure 1**). None of the samples had reduced levels of insulin (**Supplementary Figure 1**), confirming that the islets from 6-week old NOD mice already had evidence of inflammation but remained in a pre-diabetic stage without significant β-cell loss.

We next carried out lipidomics analysis of these samples, leading to the identification and quantification a total of 369, 558 and 251 lipid species in EndoC-βH1 cells, human islets and murine islets, respectively (**Figure 1A, Supplementary Tables 6-11**). A striking difference in the number of lysophospholipid species were observed comparing EndoC-βH1 cells with human or murine islets. Only 9 (2.4% of the total) lysophospholipid species were detected in EndoC-βH1 cells, compared to 89 (15.9%) and 25 (10.0%) in human and murine islets, respectively. To better understand possible lipid metabolic processes regulated in each of the models, we performed an enrichment analysis using Lipid Mini-On (Clair et al., 2019). This tool determines whether any specific lipid feature, such as head group, subclass, fatty acyl chain length or degree of unsaturation, is overrepresented among the differentially regulated lipids. In EndoC-βH1 cells, phosphatidylglycerol (PG), plasmanyl-phosphatidylethanolamine and lipids with polyunsaturated fatty acyl chains were enriched among the species regulated by the IL-1β + INF-γ treatment (**Figure 1B**). Similarly, lipids with polyunsaturated fatty acyl chains were also enriched in human islets treated with the cytokines (**Figure 1B**). In murine islets, only triacylglycerols (TG) with 58 total carbons in the fatty acyl chains were found to be significantly enriched (**Figure 1B**).

**Figure 1.**
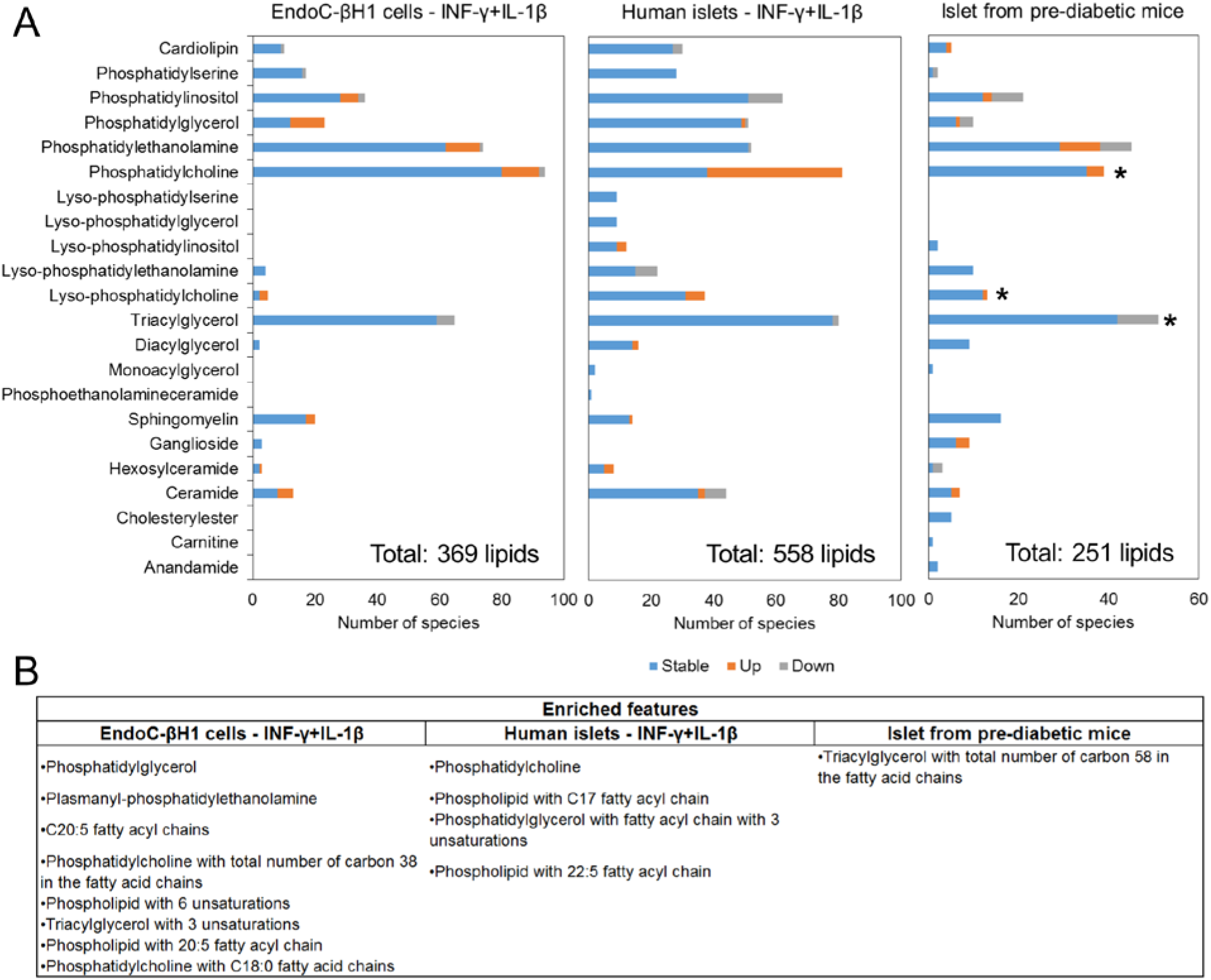
Global lipidomic analysis of 3 common models used for the study of β-cell stress in type 1 diabetes: (I) EndoC-βH1 cells exposed to IL-1β and INF-γ for 48 h (n=3), (II) human islets exposed to same cytokines for 24 h (n=10) and (III) islets from non-obese diabetic (NOD) mice in pre-diabetic stage (6 weeks of age) vs. age-matched NOR mice (n=3). Lipids were extracted and analyzed by liquid chromatography-tandem mass spectrometry. (A) Number of identified and regulated species in lipid subclass. (B) Lipid features and groups that were significantly enriched (p ≤ 0.05, Fisher’s exact test) among the differentially regulated species. *Lipid classes consistently regulated in all three samples.

We reasoned that lipid groups regulated in a coordinated/common fashion in EndoC-βH1 cells, human islets, and murine islets in response to proinflammatory cytokines were likely to be important for T1D pathogenesis (highlighted by stars in **Figure 1A**). Interestingly, LPC species, PC species with long, polyunsaturated fatty acyl chains, and TGs were commonly regulated across all three models; LPCs and PCs with long, polyunsaturated fatty acids were also upregulated, whereas TGs were consistently downregulated in all 3 sample types (**Figure 2**).

**Figure 2.**
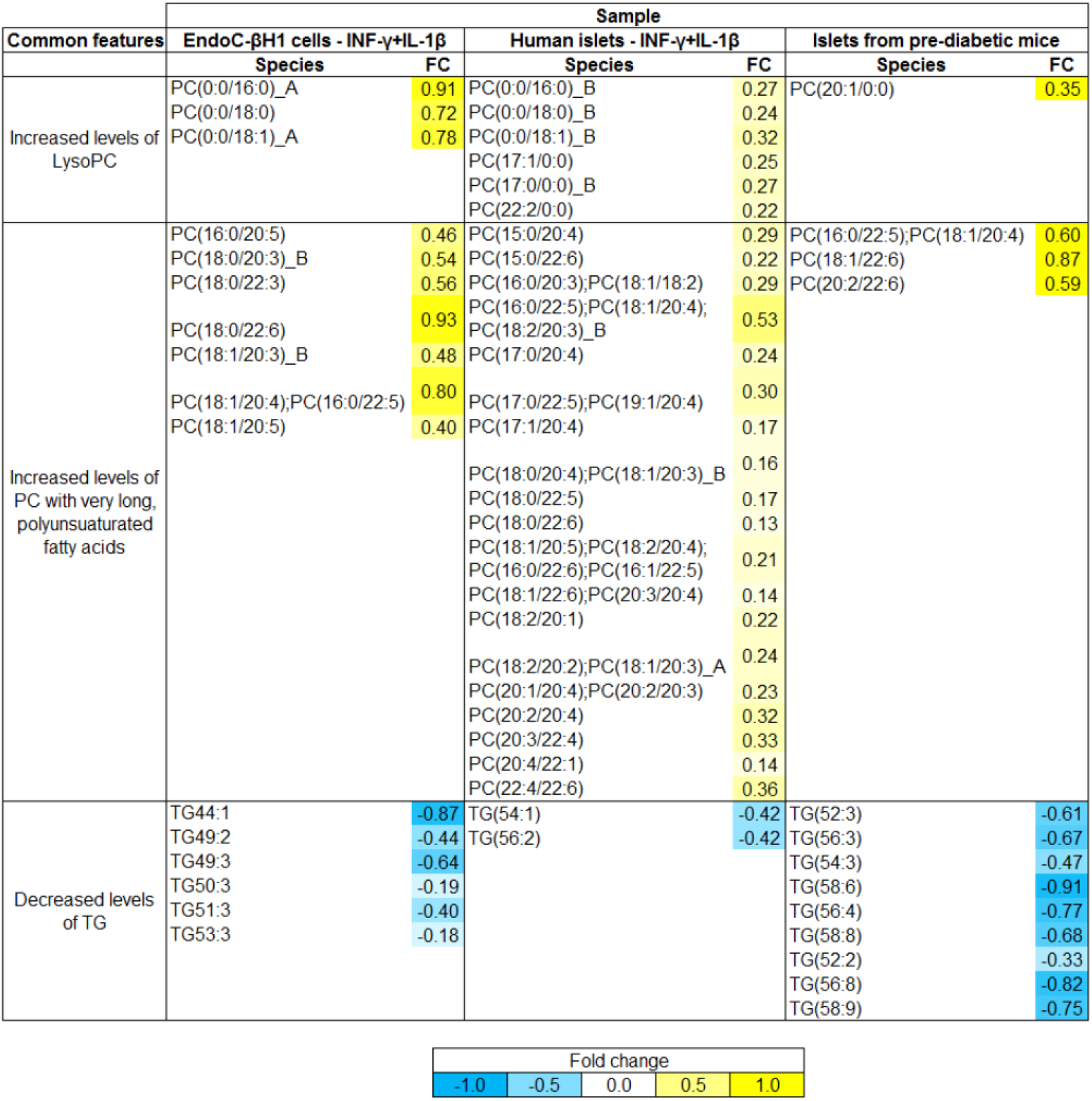
Lipid species that are consistently regulated in 3 models of β-cell stress in type 1 diabetes. Lipids from EndoC-βH1 cells treated with IL-1β and INF-γ for 48 h, human islets treated with the proinflammatory cytokines IL-1β and INF-γ for 24 h and islets from non-obese diabetic (NOD) mice in pre-diabetic stage (6 weeks of age) were extracted and analyzed by liquid chromatography tandem mass spectrometry. Each lipid species is named with the abbreviation of its class (e.g. LPC and PC) followed by the length of the fatty acid and number of double bounds (separated by colon) in parenthesis. The letters after the lipid names represent different isomers that are separated in the chromatography in alphabetical order. The relative abundance in T1D model vs. control are color-coded. Isobaric coeluting species (separated by semicolons) were co-quantified.

### Spatial distribution of coordinately regulated lipids in islets

We further surmised that coordinately regulated lipids that were particularly important for T1D pathogenesis would be preferentially localized to islets. To test this hypothesis, we determined the spatial distribution of the commonly regulated lipid species in murine pancreas using mass spectrometry imaging (MSI). MSI analysis showed that several LPC species are preferentially localized to islets of both NOD and NOR mice, including LPC(18:0) and LPC(18:1), whereas species such as LPC(18:2) and LPC(16:0) had similar abundance compared to surrounding tissue **(Figure 3A-C**). In contrast, LPC(20:4) seemed to be enriched in islets of NOR mice, while this same lipid species was depleted in islets from NOD mice (**Figure 3A-C**). As a control, the standard LPC(19:0), which is delivered with the extraction solvent, was evenly distributed throughout the tissue (**Figure 3A**). Similarly, the endogenous metabolite glycerophosphocholine (GPC) was also evenly distributed throughout the tissue (**Figure 3A**), which indicates an insignificant matrix effect (Lanekoff et al., 2014). As expected, several PC species including PC(34:1) and PC(36:2) displayed an even distribution between the islets and the surrounding tissue in both mouse strains (**Figure 3B-C**). Conversely, species with longer and polyunsaturated fatty acyl chains PC(40:5) and PC(40:6) were clearly enriched in islets from both NOR and NOD mice (**Figure 3B-C**). Unfortunately, the solvent composition used in imaging experiments did not allow us to analyze TG localization, due to inefficient extraction of this lipid class into 9:1 methanol:water mixture.

**Figure 3.**
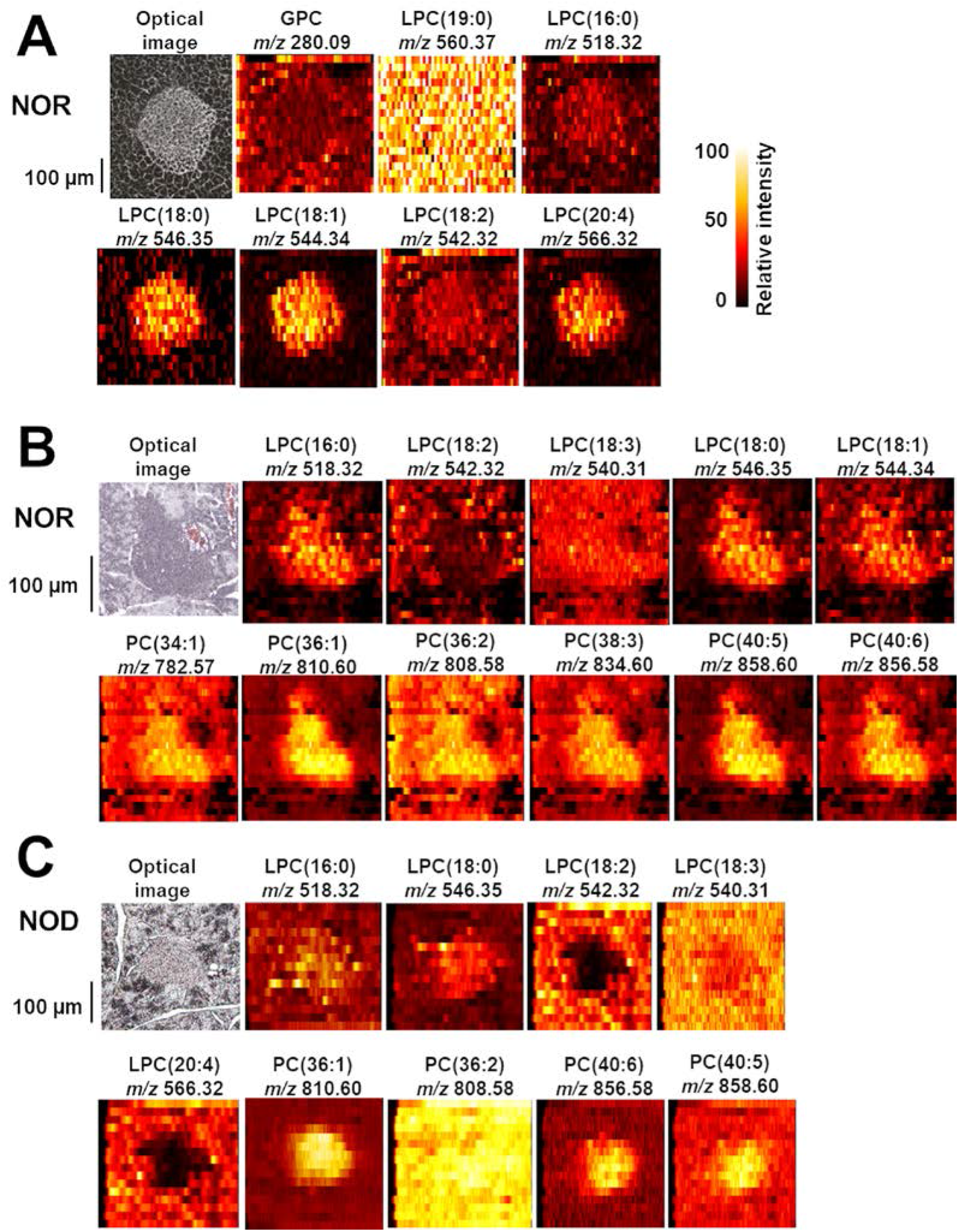
Spatial localization of lysophosphatidylcholine (LPC) and phosphatidylcholine (PC) species in pancreata from NOR (A and B) or NOD mice (C) by mass spectrometry imaging. Each image shows either the optical image or color-coded distribution of different lipid, endogenous metabolite [glycerophosphocoline (GPC)] and standard [LPC(19:0)] species. Each lipid species is named with the abbreviation of its class (e.g. LPC and PC) followed by the length of the fatty acid chains and number of double bounds (separated by colon) in parenthesis. Pancreata from 6-week old mice were collected and analyzed in independent experiments. Lipid species were identified by matching against the lipids characterized on the lipidomics analysis based on their accurate masses.

LPCs are generated by the release of fatty acid chains from phospholipids, a reaction catalyzed by phospholipases, such as the calcium-independent phospholipase A2 (iPLA2b/PLA2G6). Consistent with the localization of LPCs, *Pla2g6* was also enriched in islets as demonstrated by combining fluorescent in situ hybridization (FISH) and immunohistochemistry (**Figure 4A-B**). Taken together these data support the notion that the generation of LPCs is mediated by phospholipases, such as *Pla2g6*, which is enriched in islets compared to the surrounding tissue.

**Figure 4.**
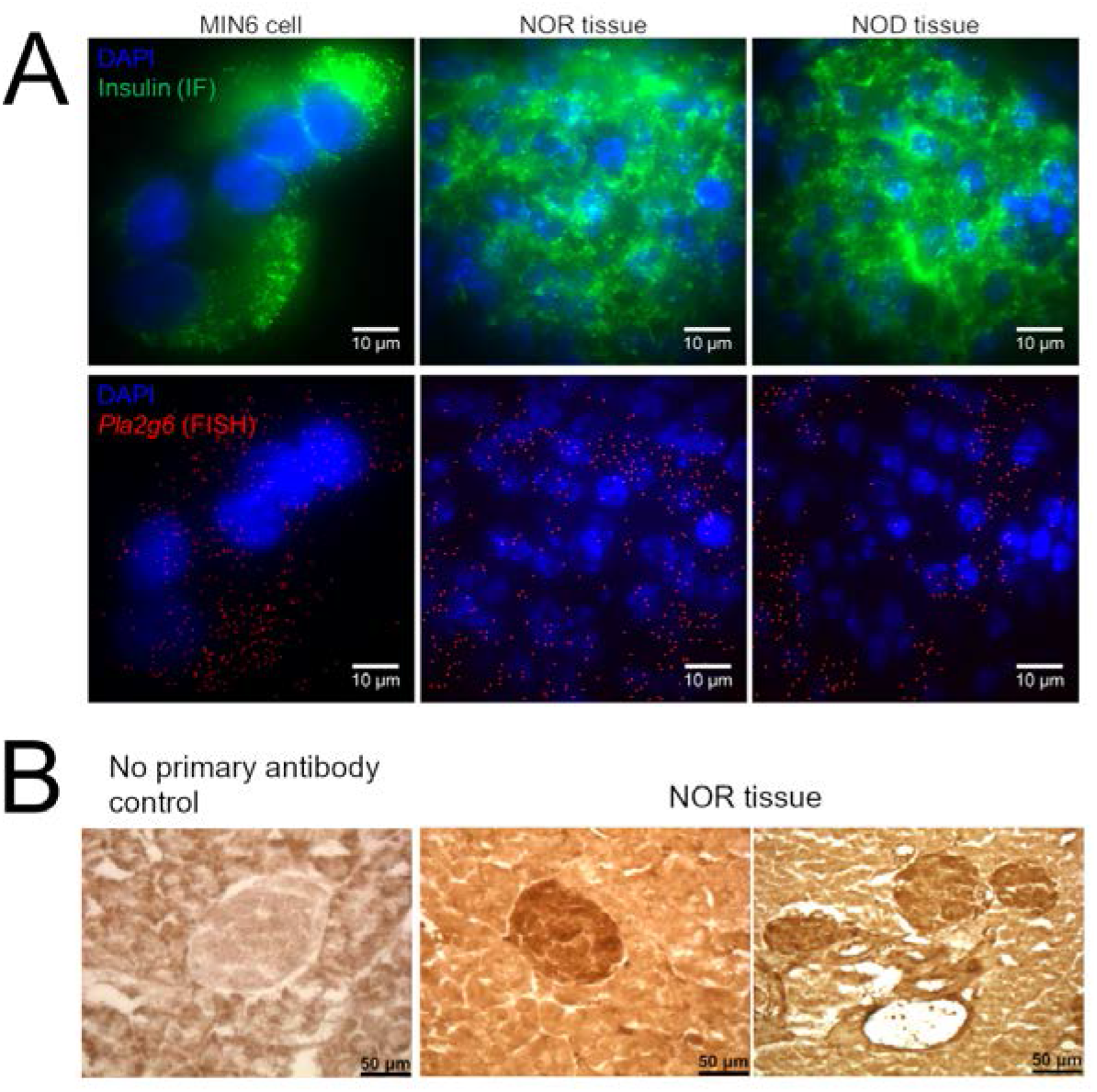
Pla2g6 distribution. (A) Fluorescence in situ hybridization (FISH) of MIN6 β cell line, and pancreata from non-obese diabetic (NOD) and non-obese diabetes resistant (NOR) mice (6 weeks of age). Cells and tissues were stained with anti-insulin antibody (green), DNA stain 4′,6-diamidino-2-phenylindole (DAPI – blue) and fluorescent-labeled antisense Pla2g6 oligonucleotide (red). (B) Immunohistochemistry (IHC) analysis of Pla2g6. Tissue was stained with biotin-conjugated anti-Pla2g6 antibodies followed by avidin-conjugated horseradish peroxidase. Localization was visualized by horseradish peroxidase-mediated oxidation and precipitation of 3,3′-diaminobenzidine (brown).

### Pla2g6-dependent cytokine regulation in MIN6 cells

To study the molecular mechanism(s) underlying phospholipase PLA2G6-mediated signaling in cytokine-exposed islets, we treated control and *Pla2g6* knockdown (KD) MIN6 β cells with IL-1β + INF-γ + TNFα for 24h, followed by LC-MS/MS-based proteomics analysis. The cytokine treatment led to changes in abundance of 1043 out of the 5212 identified and quantified proteins (**Figure 5A, Supplementary Table 12**). A function-enrichment analysis of the differentially regulated proteins revealed that 52 KEGG pathways were regulated by the cytokine treatment (**Figure 5B**). Only a small fraction of 35 proteins that were differentially regulated by cytokines was dependent on PLA2G6 (**Figure 5C, Supplementary Table 13**). Among these proteins, the expression of cathepsin Z and cathepsin B were upregulated with the cytokine treatment, but this increase in abundance was abolished in the absence of PLA2G6 (**Figures 5D-E**). Cathepsins are markers of lysosomal function and autophagy, indicating a possible link between cytokine signaling and these processes through PLA2G6. As a control, we checked the expression levels of insulin-1 and insulin-2 proteins, but they were not affected by the *Pla2g6* KD (**Figure 5F-G**). Overall, the analysis showed a strong regulation of the MIN6 β cell proteome by pro-inflammatory cytokines, of which a small subset of proteins was dependent of PLA2G6.

**Figure 5.**
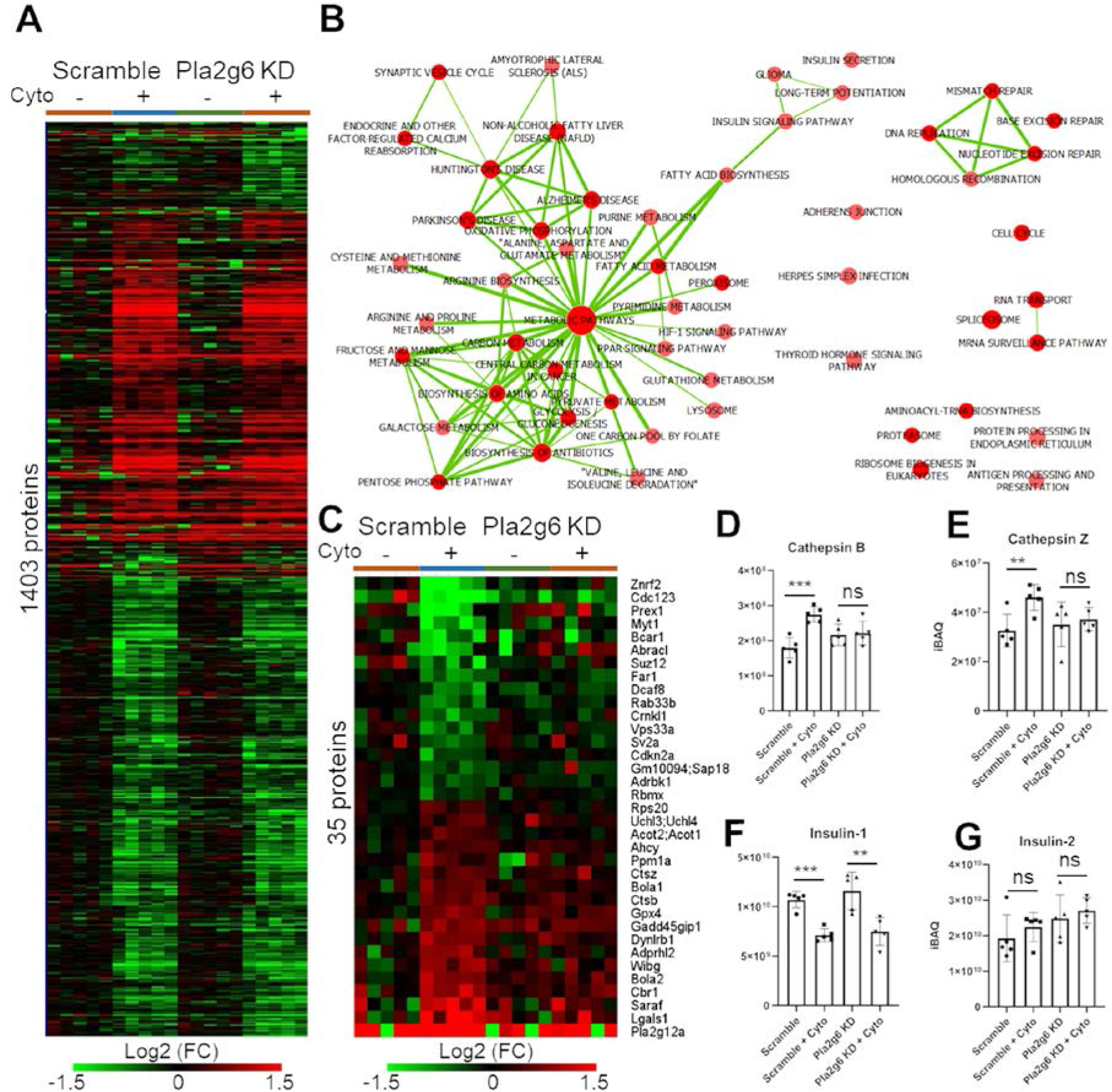
Pro-inflammatory cytokines IL-1β + IFN-γ + TNFα and Pla2g6-dependent proteome remodeling in MIN6 β cells. (A) Pro-inflammatory cytokines (Cyt)-dependent protein expression in wild-type (WT) and Pla2g6-RNAi MIN6 cells (n=5, each). (B) KEGG pathways enriched with proteins differentially abundant in IL-1β + IFN-γ-treated MIN6 cells. Pathways were grouped based on shared proteins using Enrichment Map (Merico et al., 2010). Each pathway is represented by a node and their degree of connectivity (thickness of the edges) is proportional to the number of shared proteins between the pathways. (C) Pla2g6-dependent protein abundance changes of pro-inflammatory cytokine-treated MIN6 cells. (D-G) Abundance profiles of cathepsin Z (D), cathepsin B (E), insulin-1 (F) and insulin-2 (G). Statistical test: ** p ≤0.01 and *** p≤0.001 by *t*-test considering equal distribution and variance.

### Regulation of poly(ADP)ribosylation proteins in MIN6 β cells

Among the PLA2G6-dependent cytokine-regulated proteins as judged by LC-MS proteomics was ADP-ribosyl-acceptor glycohydrolase ARH3 (ADPRHL2 gene) (**Figure 5C**). ARH3 has been shown to play a role in poly-ADP-ribose (PAR) metabolism, a factor that regulates β-cell death in mice (Andreone et al., 2012). We investigated the abundance profiles of the PAR polymerases (PARP) and hydrolases PAR glycohydrolases (PARG), MacroD1, MacroD2, terminal ADP-ribose protein glycohydrolase 1 (TARG1) and ADP-ribosyl-acceptor hydrolases (ARHs) in the proteomic analysis. PARP-1 abundance decreased approximately 35% in both *Pla2g6*-KD and control cells when treated with cytokines, whereas PARP-2 was not significantly changed (**Table 1**). Conversely, PARP-3, -9, -10, -12 and -14 were strongly upregulated (>5 fold) when cells were treated with cytokines in both wild-type and *Pla2g6*-KD cells (**Table 1**). The PAR hydrolases ARH3 and MacroD1 were also increased by 66% and 31%, respectively, in the wild-type cells treated with cytokines, but their expression was unchanged in *Pla2g6*-KD cells treated with cytokines (**Table 1**). The other PAR hydrolases were not detected in our analysis.

**Table 1:** Abundance profiles of poly(ADP)ribosylation proteins in wild-type and Pla2g6 knockdown MIN6 β cells treated with the pro-inflammatory cytokines. *iBAQ – intensity-based absolute quantification. Statistical test was done with *t*-test considering equal variation and distribution.

|  |  | Average iBAQ intensity |  |  |  | Standard deviation |  |  |  |  |  |  |  |
| --- | --- | --- | --- | --- | --- | --- | --- | --- | --- | --- | --- | --- | --- |
|  |  | Wild type |  | Pla2g6 RNAi |  | Wild type |  | Pla2g6-RNAi |  | Fold change (Treated/Control) |  | p-value (Control vs. treated) |  |
| Accession number | Protein | Control | Treated | Control | Treated | Control | Treated | Control | Treated | Wild type | Pla2g6 RNAi | Wild type | Pla2g6 RNAi |
| Q921K2 | PARP-1 | 160541298 | 102181010 | 162185126 | 98199854 | 14415018 | 13769483 | 11315442 | 11911652 | 0.64 | 0.61 | 0.000 | 0.000 |
| O88554 | PARP-2 | 2320187 | 1886698 | 2701832 | 1904552 | 773539 | 900914 | 606622 | 619003 | 0.81 | 0.70 | 0.438 | 0.074 |
| Q8CFB8 | PARP-3 | 0 | 2440297 | 129711 | 2796473 | 0 | 1950741 | 290043 | 1320074 | ∞ | 21.56 | 0.023 | 0.002 |
| Q8CAS9 | PARP-9 | 1027221 | 39521623 | 1758362 | 44078979 | 833995 | 6228610 | 1232660 | 4239980 | 38.47 | 25.07 | 0.000 | 0.000 |
| Q8CIE4 | PARP-10 | 100029 | 4173449 | 93000 | 4561784 | 223672 | 1064209 | 207955 | 1039258 | 41.72 | 49.05 | 0.000 | 0.000 |
| Q8BZ20 | PARP-12 | 613429 | 3541763 | 861360 | 6368375 | 261033 | 682862 | 471954 | 1427301 | 5.77 | 7.39 | 0.000 | 0.000 |
| Q2EMV9 | PARP-14 | 616310 | 26623139 | 769444 | 27348827 | 84838 | 2863548 | 321341 | 1432066 | 43.20 | 35.54 | 0.000 | 0.000 |
| Q8CG72 | ARH3 | 11541195 | 19182811 | 11421375 | 12104702 | 3610258 | 2655778 | 3105854 | 1940956 | 1.66 | 1.06 | 0.005 | 0.688 |
| Q922B1 | MacroD1 | 16322818 | 21400784 | 19062601 | 17347089 | 3285980 | 2286317 | 3446722 | 3884448 | 1.31 | 0.91 | 0.022 | 0.481 |

To further investigate the role of ARH3 in β-cell stress, we tested the effects of ARH3 KD in cytokine-mediated apoptosis by performing western blots of the apoptotic marker cleaved caspase 3. The KD reduced ARH3 protein levels by 65-72% (**Figure 6A-B**). The cytokine cocktail IL-1β + INF-γ + TNFα increased the expression level of cleaved caspase 3 by 8.8 fold in the scramble RNAi cells as compared to the non-cytokine treated control group (**Figure 6A and 6C**). In ARH3 KD cells, the levels of cleaved caspase 3 were further increased by 15.4 fold (**Figure 6A and 6C**). Of note, ARH3 KD already had a higher basal level of cleaved caspase 3 (**Figure 6A and 6C**).

**Figure 6.**
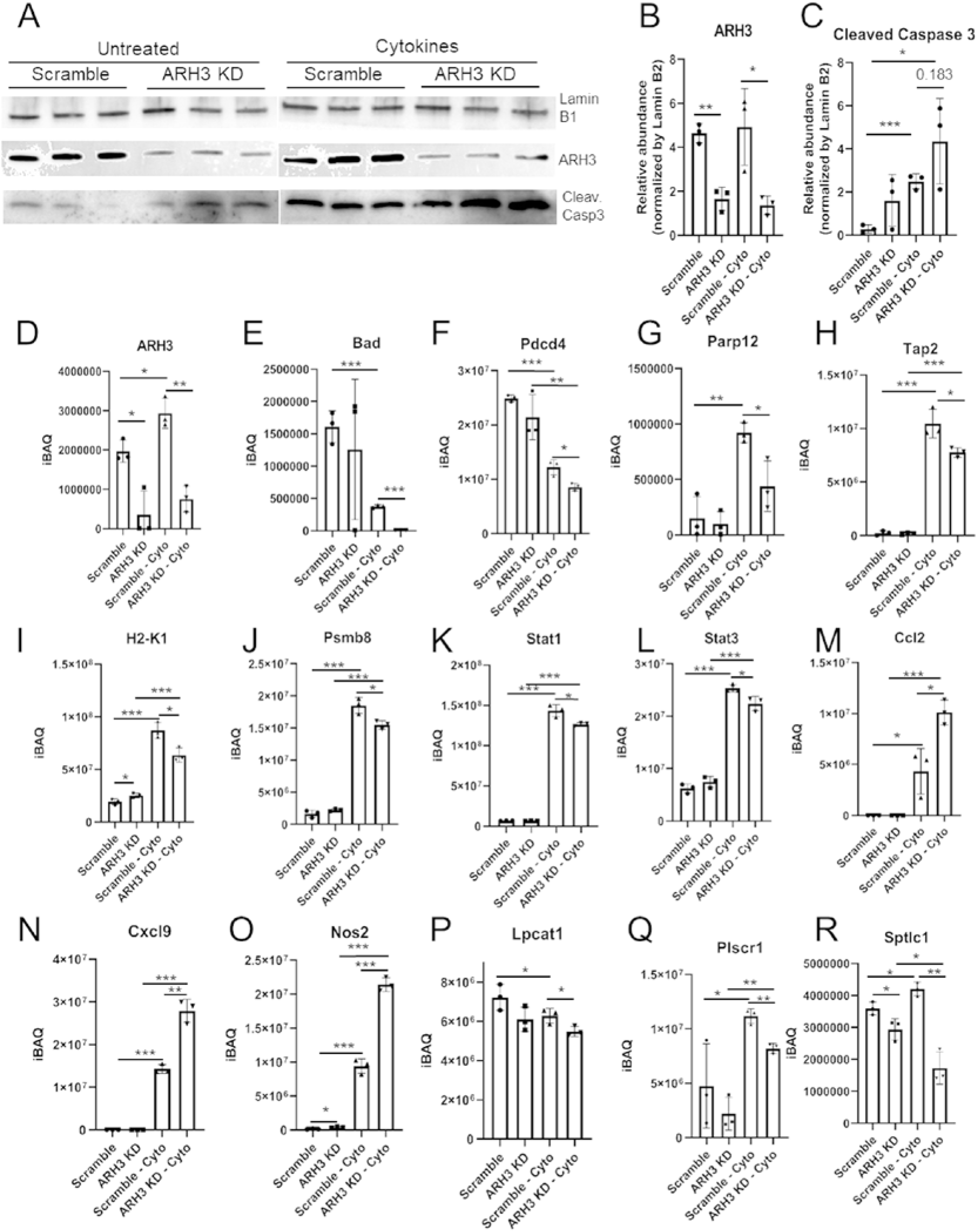
The role of ADP-ribosy-acceptor glycohydrolase ARH3 in cytokine-signaling of MIN 6 β cells. MIN6 cells were transfected with ARH3 RNAi or scrambled oligonucleotides and treated with 100 ng/mL IFN-γ, 10 ng/mL TNF-α, and 5 ng/mL IL-1β for 24 h and then analyzed by western blot and proteomics analysis. (A) Western blot analysis of ARH3 KD cells treated with cytokines. (B-C) Quantification of the levels of ARH3 (B) and cleaved caspase 3 (C) bands. To ensure the reproducibility we performed experiment multiple times. The data is representative of 4 independent experiments and 6 replicates total with similar results. (D-R) Quantification of selected proteins from the proteomics analysis (**Supplementary Tables 14-16**): (D) ARH3, (E) Bad: Bcl2-associated agonist of cell death, (F) Pdcd4: Programmed cell death protein 4, (G) Parp12: Poly [ADP-ribose] polymerase 12, (H) Tap2: Antigen peptide transporter 2, (I) H2-K1: H-2 class I histocompatibility antigen K-B alpha chain, (J) Psmb8: Proteasome subunit beta type-8, (K) Stat1: Signal transducer and activator of transcription 1, (L) Stat3: Signal transducer and activator of transcription 3, (M) Ccl2: C-C motif chemokine 2, (N) Cxcl9: C-X-C motif chemokine 9, (O) Nos2: Inducible nitric oxide synthase, (P) Lpcat1: Lysophosphatidylcholine acyltransferase 1, (Q) Plscr1: Phospholipid scramblase 1, (R) Sptlc1: Serine palmitoyltransferase 1. Statistical test: * p ≤0.05, ** p ≤0.01 and *** p≤0.001 by *t*-test considering equal distribution and variance.

To gain more mechanistic insights, we performed proteomic analysis of the same samples (**Supplementary Tables 14-15**), which confirmed the reduction in ARH3 expression in the KD cells (**Figure 6D**). The levels of the pro-apoptotic Bcl-2 protein Bad and programmed cell death protein 4 (Pdcd4) were reduced by the cytokine treatment and the ARH3 KD further reduced their levels (**Figure 6E-F**). Conversely, Parp12 levels increased with the cytokine treatment, but the ARH3 KD reduced this upregulation (**Figure 6G**). Parp12 was the only PARP regulated by cytokines in a PLA2G6-dependent fashion (**Table 1**), and since it was regulated in opposite direction in ARH3 KD it could represent a negative feedback mechanism. A function-enrichment analysis of the proteomics data showed that pathways such as antigen processing and presentation, and chemokine signaling, were regulated by cytokines and ARH3 (**Supplementary Table 16**). In terms of antigen processing and presentation, the antigen transporter protein Tap2, major histocompatibility complex I subunit H2-K1 and immunoproteasome subunit Psmb8 were consistently reduced in ARH3 KD cells as compared to scramble RNAi when cells were treated with cytokines (**Figure 6H-J**). Among the chemokine-regulating proteins, the transcription regulators Stat1 and Stat3 had both a minor (12%) reduction in ARH3 KD cells treated with cytokines as compared to the scramble RNAi cells (**Figure 6K-L**). Conversely, the ARH3 KD enhanced the expression of some cytokine-induced genes, such as the chemokines Ccl2 and Cxcl9, and nitric oxide synthase Nos2 (**Figure 6M-O**). We also observed that ARH3 KD reduced the abundance of lipid-related proteins, such as lysophosphatidylcholine acyltransferase Lpcat1, phospholipid scrambalase Plscr1 and serine palmitoyltransferase Sptlc1 (**Figure 6P-R**), suggesting a role of ARH3 in regulating lipid metabolism. Together, these data show a strong regulation of the poly(ADP)ribosylation machinery by cytokines in MIN6 β cells and suggest that ARH3 is further regulated by PLA2G6, leading to β cell protection from apoptosis through a complex regulatory network.

## DISCUSSION

We investigated the remodeling of islet and β cell lipid composition in 3 models of T1D. Since phenotypes found in model systems do not always reflect the physiology of disease, we performed lipidomic analysis across three different models type 1 diabetes: EndoC-βH1 cells exposed to the cytokines IL-1β + INF-γ, human islets exposed to the same cytokines and islets from non-obese diabetic (NOD) mice at the pre-diabetic stage. This analysis identified consistent changes in composition of specific classes of lipids: down regulation of TGs and upregulation of LPCs and PCs with long unsaturated fatty acid chains. TGs have been shown to be down-regulated in rat β cells by the pro-inflammatory cytokine combination IL-1β + INF-γ + TNFα, likely to accommodate the resulting increased energetic demands (Kiely et al., 2007). TG metabolism has also been associated with cellular protection; it has been shown that GDF15 impairs inflammation-induced damage by systemic release of liver TGs (Luan et al., 2019). Consistently, TG-carrying protein apolipoprotein CIII reduces apoptosis in rodent β cells treated with IL-1β + INF-γ (Storling et al., 2011). Also, increases in cellular TG content has been associated with protection of rat β cells against the cytotoxic effects of saturated free fatty acids (Cnop et al., 2001).

Our three different models of T1D also showed a consistent upregulation in the levels of PCs with long polyunsaturated fatty acid chains. Like TGs, polyunsaturated fatty acids have been associated with protection of β cells against pro-inflammatory cytokines (Wei et al., 2010). An omega 3 polyunsaturated fatty acid diet reduces the incidence of diabetes in NOD mice (Bi et al., 2017). Polyunsaturated fatty acids are sources for immunomediators such as prostaglandins, leukotrienes and thromboxanes. While some of these immunomediators are anti-inflammatory, others induce inflammation and apoptosis. For instance, the processing of the polyunsaturated fatty acid arachidonic acid into 12-HETE by 12-lipoxygenase has been shown to lead to β-cell death (Tersey et al., 2015).

Polyunsaturated fatty acids are often associated with phospholipids but can be released from them by phospholipases. Multiple phospholipases may contribute to this process as we detected dozens of lipases in the proteomic analysis (**Supplementary Tables 4, 12 and 14**). We focused in the inducible phospholipase A2 beta (iPLA2b/PLA2G6) as it has been reported to regulated by cytokines in islets, releasing of polyunsaturated fatty acids and LPCs (Lei et al., 2014; Lei et al., 2010). This is in line with our results showing an accordant increase of LPCs in each of the T1D models. We also showed that several LPCs are enriched in islets, which is consistent with the observation that PLA2G6 expression is higher in islets compared to surrounding tissue. Our previous proteomics analysis showed that PLA2G6 is also expressed in human islets (Nakayasu et al., 2020a), while a previous report showed that its transcript is higher in rat β cells compared to α cells (Ma et al., 1998). PLA2G6 has been shown to activate apoptosis via downstream activation of neutral sphingomyelinase and formation of ceramides, leading to mitochondrial membrane decompensation (Lei et al., 2014). Pharmacological inhibition of PLA2G6 protects NOD mice from developing diabetes (Bone et al., 2015). PLA2G6 can also induce alternative splicing and reduce the levels of the antiapoptotic protein Bcl-x(L) via neutral sphingomyelinase activation, ceramide production and ER stress (Barbour et al., 2015). We found upregulated species of ceramides in all three different models of T1D used (**Figure 1**). However, the IL-1β + INF-γ treatment also downregulated the concentration of several ceramide species in human islets but not in the other 2 studied models (**Figure 1**), suggesting that additional processes might occur in human islets. PLA2G6 was previously shown to regulate autophagy and apoptosis in murine islets treated with the ER stressor, thapsigargin (Lei et al., 2013). This observation agrees with our proteomics data, which shows that PLA2G6 regulates lysosomal trafficking and autophagy markers cathepsin B and Z. Of note, another cathepsin, cathepsin H, is a candidate gene for type 1 diabetes with a key role for β-cell function and survival (Floyel et al., 2014) and crinosomes, a cell structure that plays a key role for the generation of autoimmune peptides in diabetes prone NOD-mice (Wan et al., 2020), are rich in cathepsins.

Here we show that PLA2G6 also regulates antiapoptotic signals by inducing the expression of the ADP-ribosyl-acceptor glycohydrolase ARH3. ARH3 hydrolyzes PAR from serine residues, counterbalancing the pro-apoptotic activity induced by ADP-ribose polymerization in oxidative stress (Abplanalp et al., 2017; Fontana et al., 2017; Mashimo et al., 2019). The molecular regulation of ARH3 is virtually unknown. PLA2 products, such as LPC and arachidonic acid induce signal transduction that activates the transcription factors NF-κB and AP-1, respectively (Becuwe et al., 2003; Carneiro et al., 2013). However, their role in regulating ARH3 is being pursued in ongoing studies.

In murine islets, PARP-1 has been shown to be involved in cytokine mediated β-cell death and PARP-1 deletion has been shown to protect against streptozotocin-mediated diabetes (Andreone et al., 2012; Burkart et al., 1999; Masutani et al., 1999). Our data show that this scenario might be more complex as PARP-3, -9, -10, -12 and -14 are strongly upregulated in cells treated with the cytokines IL-1β + INF-γ + TNFα, but their role in β-cell death still needs to be investigated. Of these PARPs, we found that PARP-12 expression was induced by cytokines. However, while the PLAG2G6 KD further increased the PARP-12 levels, the ARH3 KD reduced its abundance, which may represent a negative feedback mechanism. ARH3 KD further the cytokine-mediated downregulation of the apoptosis related proteins Bad and Pdcd4. ARH3 KD decreased the expression levels of proteins involved in antigen processing and presentation, a process tightly related to the development of autoimmunity and T1D (DiMeglio et al., 2018). Conversely, ARH3 KD increased the abundance of the pro-inflammatory cytokine-induced proteins, such as Nos2 (Suarez-Pinzon et al., 2001), and the chemokines Ccl2 and Cxcl9 (Karin et al., 2016), while decreasing the expression of lysophosphatidylcholine acetyltransferase Lpcat1, phospholipid scrambalase Plscr1 and serine palmitoyltransferase Sptlc1. The regulation of Lpcat1, Plscr1 and Sptlc1 proteins by ARH3 suggests that it may contribute to maintaining lipid homeostasis. Intriguingly, all these proteins have been previously shown to have pro- or anti-apoptotic functions and/or to contribute to T1D development (Akagi et al., 2016; Karin et al., 2016; Ramos-Rodriguez et al., 2019; Sivagnanam et al., 2017; Suarez-Pinzon et al., 2001; Taouji et al., 2013). Therefore, our data suggest that ARH3 promotes β-cell survival by balancing pro- and anti-apoptotic factors through a very complex regulatory network (**Figure 7**).

**Figure 7.**
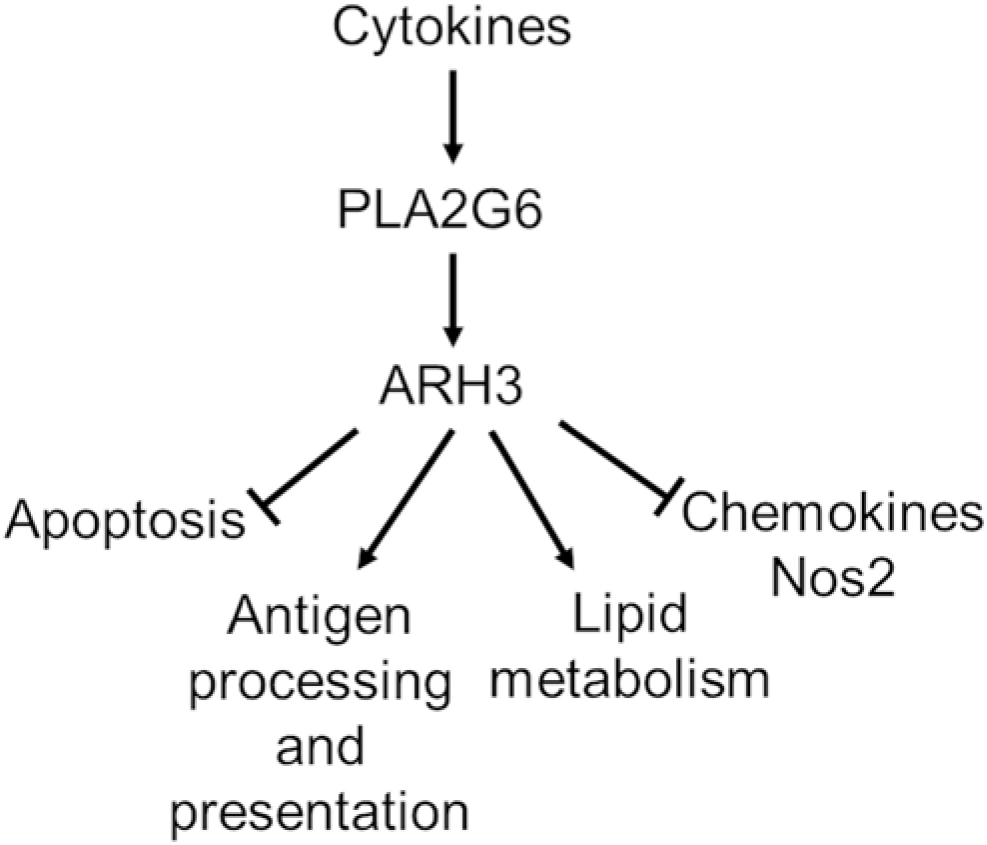
Model of cytokine signaling regulated by phospholipase PLA2G6 and ADP-ribosyl-acceptor glycohydrolase ARH3. Cytokines activates PLA2G6 that induces an upregulation of ARH3. ARH3 in turn suppresses apoptosis and cytokine-induced genes, such as chemokines and nitric oxide synthase (Nos2). ARH3 also enhances the expression of genes related to antigen presentation and lipid metabolic proteins.

Overall, our data showed a consistent regulation of the lipid metabolism in 3 different models of insulitis. Additionally, we demonstrated that phospholipase Pla2g6 signaling leads to the upregulation of ARH3 which triggers a complex regulatory network that provides a feedback mechanism that reduces apoptosis.

## Acknowledgements

The authors thank the NIDDK-supported Integrated Islet Distribution Program (IIDP) for providing the human islets used in the study. Work was performed in the Environmental Molecular Sciences Laboratory, a U.S. Department of Energy (DOE) national scientific user facility at Pacific Northwest National Laboratory (PNNL) in Richland, WA. Battelle operates PNNL for the DOE under contract DE-AC05-76RLO01830. A non-peer reviewed, preprint version of this work was uploaded into Biorxiv under DOI: https://doi.org/10.1101/2020.03.23.004481.

## Authors’ contributions

E.S.N., L.S., C.A. conceived the study and participated in the study design. E.S.N., M.G., J.K., D.S., C.D., R.Y., Y.C., C.N., F.S., and J.J.M. performed the experiments. All the authors performed the data analysis. E.S.N., R.Y., Y.C., B.J.W.R., J.L., L.S. and C.A. wrote the manuscript. All authors read, revised and approved the final manuscript for publication.

## Data and Resource Availability

Proteomics data were deposited into Pride repository (www.ebi.ac.uk/pride) under accession number PXD017863, PXD021501 and PXD021475. Lipidomics data were deposited into Massive repository (http://massive.ucsd.edu/) under accession number MSV000086174.

Pla2g6-dependent cytokine signaling in MIN6 cells

Project number: PXD017863

Reviewer account details:

Username

Password: B4o1VcPd

ARH3-regulated cytokine signaling in MIN6 cells

Project accession: PXD021501

Reviewer account details:

Username

Password: NbhKnd9N

Proteomics analysis of islets of 6-week old NOR and NOD mice

Project accession: PXD021475

Reviewer account details:

Username

Password: tKFfLzrh

Lipidomics of 3 models of insulitis, beta-cell stress and type 1 diabetes development

Project accession: MSV000086174

Reviewer account details:

Username: MSV000086174_reviewer

Password: Islets3663

## Legends

**Supplementary Figure 1.**
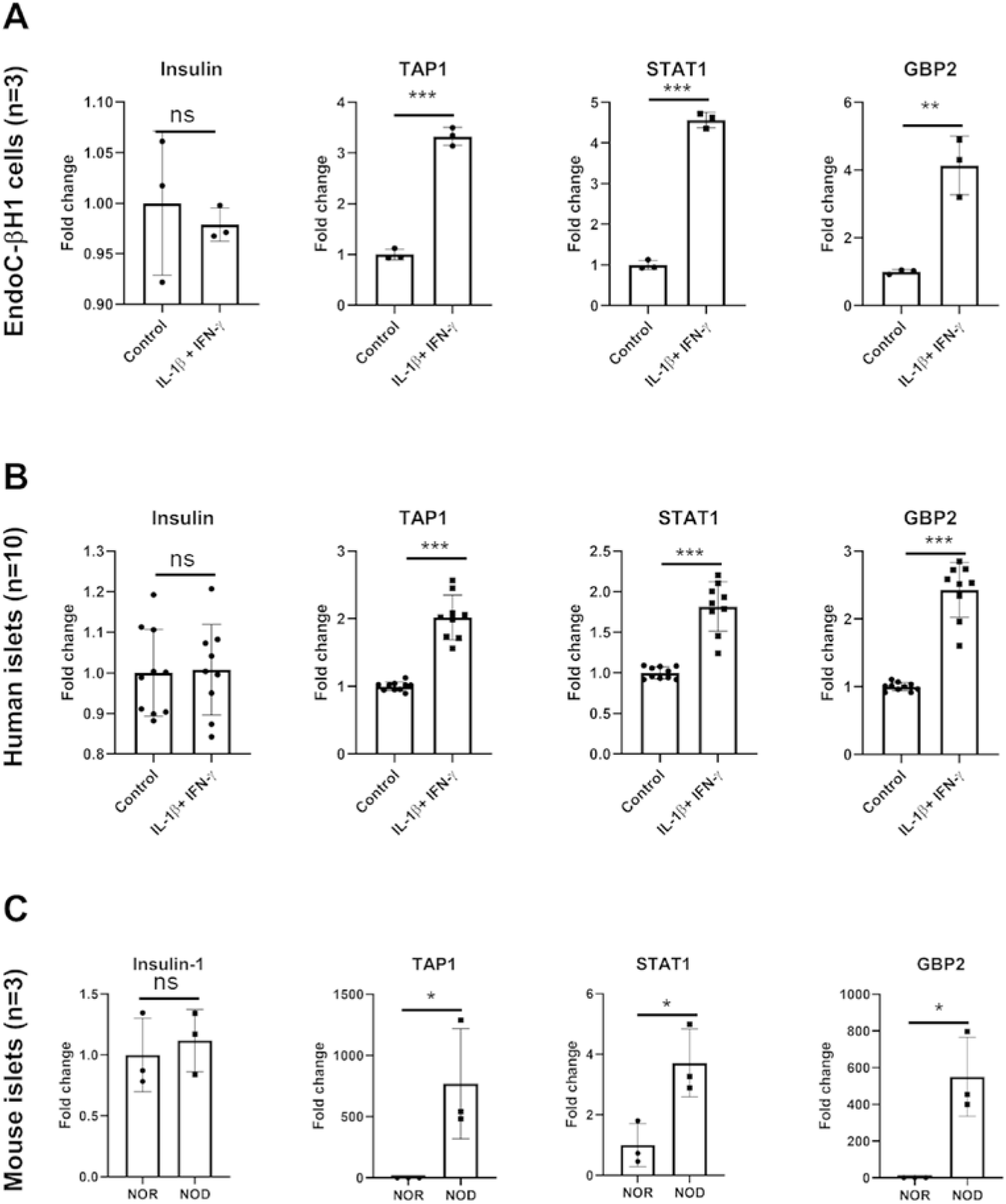
Abundance of selected proteins from proteomics analysis of 3 common models used for the study of β-cell stress in type 1 diabetes: (A) EndoC-βH1 cells exposed to IL-1β and INF-γ for 48 h (n=3) (Ramos-Rodriguez et al., 2019), (B) human islets exposed to same cytokines for 24 h (n=10) (Nakayasu et al., 2020b) and (C) islets from non-obese diabetic (NOD) mice in pre-diabetic stage (6 weeks of age) vs. age-matched NOR mice (n=3) (**Supplementary Tables 4-5**). Abbreviations: GBP2: interferon-induced guanylated-binding protein 2, Stat1: signal transducer and activator of transcription 1, TAP1: antigen peptide transporter 1. Statistical test: ** p ≤0.01 and *** p≤0.001 by *t*-test considering equal distribution and variance.

